# A comparative dataset on population genetics, traits, and distributions for nineteen Caribbean tree species

**DOI:** 10.64898/2026.03.21.713382

**Authors:** Laura Moro, Pascal Milesi, Betsaida Cabrera García, Teodoro Clase, Francesc Borràs Sayas, Eleanor Gibney, Yommi Piña, María Uriarte, Robert Muscarella

## Abstract

Although genetic diversity is a fundamental component of biodiversity, we lack data for a majority of species, particularly in biodiversity hotspots such as tropical forests. We present a comparative genetic dataset of 19 tropical tree species (including one palm) from the Caribbean island of Puerto Rico and neighboring islands (Hispañola and the US Virgin Islands). Using a reduced-representation sequencing technique (SLAF-seq), we identified species-specific single-nucleotide polymorphism (SNP) datasets with 24,413 to 433,637 high-quality SNPs per species. The focal species represent a range of life-history and climate associations, which may be relevant to their genetic structure. Therefore, we also include complementary information on species functional traits (wood density, leaf thickness, specific leaf area, maximum height, and seed dry mass), as well as geographic distributions and climatic associations from species distribution models.

## BACKGROUND & SUMMARY

Genetic diversity is a constitutive and yet often overlooked dimension of biodiversity. The study of genetic diversity is fundamental for quantifying the legacies and long-term effects of habitat loss and gain, assessing population status, and predicting population persistence over time ^1^. However, patterns of genetic variation have been examined in only a small fraction of species worldwide. Despite harboring exceptionally high levels of biodiversity, only a small percentage of tropical forest species have published genetic data available, underscoring the gap between biodiversity richness and available molecular resources ^2^. This data deficiency is particularly concerning in light of the escalating vulnerability of tropical forest ecosystems to anthropogenic pressures, including land-use and climate change ^3–5^.

The gap in available genetic data is especially pronounced in hyper-diverse tropical islands, such as the Antilles. These fragile ecosystems, which have experienced some of the most dramatic habitat changes over the past two decades ^6,7^, are severely underrepresented in genetic studies. Island species often exhibit lower levels of genetic diversity due to geographic isolation and small population sizes ^8^. Alarmingly, global studies have reported a ∼6% decline in genetic diversity since the industrial revolution, with island species experiencing a disproportionately high average loss of ∼28% ^9^. This highlights the urgent need to assess and monitor genetic diversity, particularly in vulnerable island ecosystems, where species may already be at greater risk of extinction.

Reduced-representation sequencing (RRS) techniques, including RAD-seq and SLAF-seq ^7^, have expanded access to high-throughput genetic data, particularly for non-model species for which reference genomes are unavailable. RRS has become a widely adopted method for ecological and evolutionary studies because it enables the genotyping of thousands to millions of SNPs in many individuals with a limited budget ^8,9^. RRS opens the door to comparative genetic studies of forest trees, such as exploring how genetic patterns are linked to life-history traits, phylogeography, and historical vegetation dynamics ^10–13^.

Complementary information on species climatic associations and life-history traits can provide additional insights into comparative genetics studies. For example, some studies have suggested that differences among species’ genetic diversity and structure are related to their dispersal abilities and successional associations ^13^. Species-specific habitat associations (e.g., climate, soils) can also help interpret patterns of genetic structure by giving insight into potential barriers to gene flow ^11^.

We present a comparative genomic dataset for 19 tree species (including one palm) from the Caribbean islands of Puerto Rico, Hispañola, and the US Virgin Islands. Using a sampling strategy designed to span species distributions across environmental gradients and RRS sequencing, we obtained population-level genomic data from a total of 790 individuals across the study species. To contextualize genetic diversity within Puerto Rico, we also collected samples from neighboring islands of Hispañola (Dominican Republic) and the US Virgin Islands. There is currently little to no genetic data available for these species, nor are there any reference genomes.

We selected the focal species to represent a wide variety of life-history traits and climatic associations (Table 1). For each species, we also provide a set of georeferenced occurrence records, results from a species distribution model (SDM) representing habitat suitability in Puerto Rico, and mean values of reproductive and vegetative life-history traits. This dataset enables analyses that explore the ecology, evolution, and biogeography of tropical trees, while also alleviating the lack of genetic data available for tropical trees, especially in species-rich island systems. Ultimately, the data provided here may help to guide species conservation efforts.

**Table 1.**
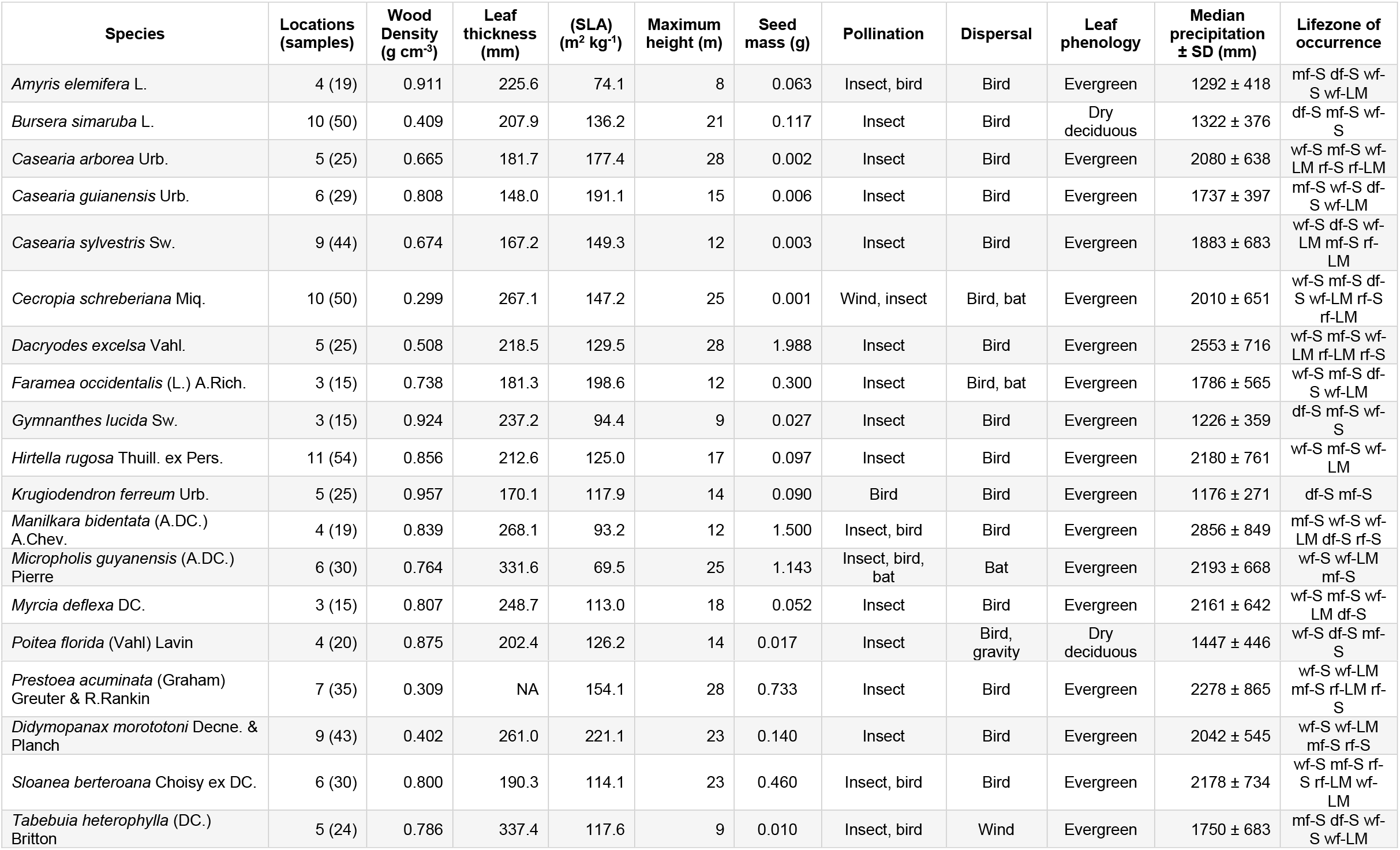
Sample size for genetic data, species mean values of traits, and climate associations for the 19 species in this dataset. Pollination and dispersal information is based on morphology and observations reported in. Precipitation values refer to the mean annual precipitation across the occurrence locations for each species. Abbreviations of Holdridge life zones correspond to the legend in Figure 1. Seed mass comes from ^28,29^ ; pollination and seed dispersal categories come from information in ^15^.

## METHODS

### Study area

The majority of our samples (72%) come from Puerto Rico, a Caribbean island in the Greater Antilles with a total land area of ∼9,000 km^2^ and a maximum elevation of 1,338 m a.s.l. (Figure 1). The island encompasses broad climatic gradients: mean annual precipitation ranges from about 900 to >4,500 mm yr^-1^ and there is also a high degree of edaphic heterogeneity in Puerto Rico, with the most common soil types being derived from volcanic and limestone parent materials ^14^. In order to contextualise the magnitude and patterns of genetic diversity within Puerto Rico, we collected a smaller number of samples from several sites on the island of Hispañola (Dominican Republic), as well as several sites on the islands of St. John and St. Thomas in the US Virgin Islands. Our collecting sites in Hispañola varied in environmental conditions from wet to dry forests while sites in the US Virgin Islands were subtropical moist and dry forests.

**Figure 1.**
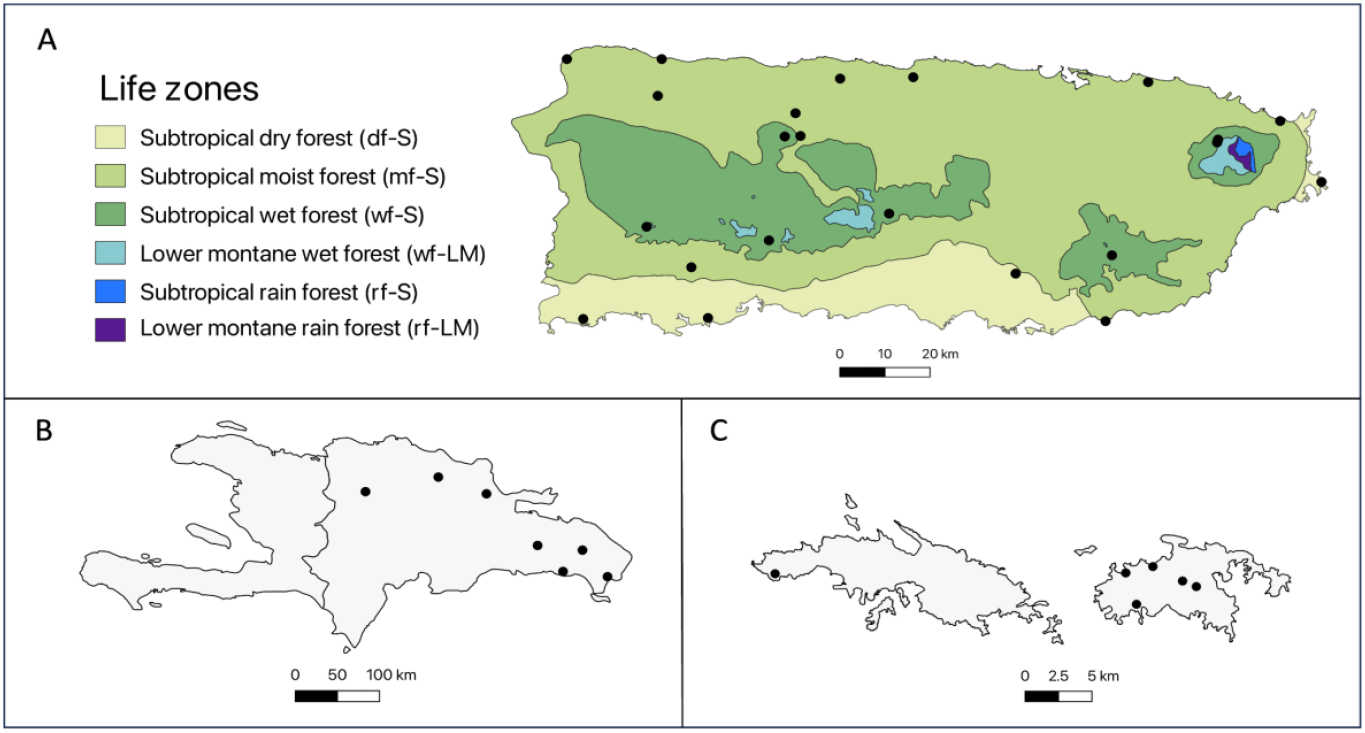
Maps of sampling locations. (A) Sample locations in Puerto Rico with Holdridge life zones^16^. (B) Sample locations in Hispañola (Dominican Republic) and (C) the US Virgin Islands.

### Sample collection

Between 2022 and 2024, we collected leaf samples from 790 individuals of 19 tree species on the Islands of Puerto Rico (N=565 for 19 species), Hispañola (N=105 for 13 species), and the US Virgin Islands (N=120 for 16 species) (Figure 1). We identified species using dichotomous keys ^15^, herbarium specimens, and expert knowledge. In Puerto Rico, the sampling extended across the known geographic range of each species, and included forests of different ages, with 4–12 locations sampled per species (Table 1). Sampling in Hispañola and the US Virgin Islands was more sparse (1–2 locations per species), as the aim was to include neighboring reference populations to contextualize genetic diversity within Puerto Rico (Figure 1).

At each site, we collected leaves from 5–10 individuals (for each species present in the site), at least 30 m apart from one another. We air-dried the samples and preserved them in silica gel until DNA extraction. For each sample, we recorded geographical coordinates with a handheld GPS and collected an herbarium voucher. The vouchers are stored at Uppsala University, Uppsala, Sweden. Samples in Puerto Rico were collected under permit 2022-IC-003 from the Departamento de Recursos Naturales y Ambientales (DNRA). Samples from the Hispañola (Dominican Republic) were collected under permit DG-22 from the Ministerio de Medio Ambiente y Recursos Naturales. Samples from the US Virgin Islands were collected under permit DFW24018U from the Department of Planning and Natural Resources Division of Fish and Wildlife.

### Sequencing

We extracted DNA from the dry leaf samples using the Qiagen DNeasy Plant Mini Kit, Qiagen^®^. We modified the manufacturer’s protocol by increasing the incubation time to 60 minutes and adding a dry centrifuge step before the two elution to obtain higher DNA quality and yield. When necessary, and in order to obtain the DNA quantity, concentration, and purity required for sequencing, we purified the extracts using the Genomic DNA Clean & Concentrator-10, Zimo Research^®^, and increased the DNA concentration by evaporation using a speed vacuum (see Supplementary Table 1 for quality metrics).

Because no reference genomes exist for our focal species, we genotyped individuals using high-resolution specific-locus amplified fragment sequencing (SLAF-seq). SLAF-seq is an RRS sequencing strategy for large-scale *de novo* SNP discovery and genotyping optimized for avoiding repetitive regions and maximizing uniform distribution across the genome ^17^. We first performed a pilot sequencing experiment to select the best combination of restriction enzymes. For each species, one sample was sequenced three times using the same restriction enzyme combination (RsaI and HaeIII) but different insert sizes (gel cutting ranges 400-480 bp, 450-500 bp, 500-600 bp). Based on the results of this pilot study, we sequenced all species with HaeIII+RsaI enzymes with the same insert size range of 364-464 bp (Illumina NovaSeq platform, pair-end 150 bp reads, targeting 100,000 tags per species and ∼10x read depth of coverage). Development and sequencing work was performed at Biomarker Technology (BMK) GmbH, Germany (www.bmkgene.com).

### Species distribution models

We built species distribution models (SDMs) for each species with Maxent 3.4.1 ^18^ using the R Package ENMeval ^19,20^. We used a dataset of 16,146 occurrence records of all Puerto Rican trees comprised of observations from GBIF(www.gbif.org), four herbaria (UPRRP, MAPR, NY, US), georeferenced data points from previously published studies^21^, as well as our own observations. The number of georeferenced occurrence points for our focal species ranged from 47–240 (median 91 ±SD 57). As explanatory variables in the SDMs, we used four abiotic climatic variables: log-transformed precipitation in the driest month (mm), an index of precipitation seasonality computed as in ^22^, temperature in the hottest month (°C), and a categorical map of geological substrate^23^. The temperature and precipitation data (climatological averages from 1963-1995) at 450 m resolution were downloaded from the PRISM Climate Group (http://prism.oregonstate.edu).

To fit the SDMs, we used the target-group background approach, which helps account for the spatial sampling bias in the occurrence records ^24^. Specifically, when building an SDM for each of the focal species, all records from all other species were used as background points. We fit models using all pairwise comparisons of five regularization multipliers (1, 2, 3, 4, 5) and four feature class combinations (“L”, “LQ’’, “LQH’’, “H”). We used spatially-blocked cross-validation to evaluate the performance of SDMs. Specifically, we partitioned the data using the checkerboard 2 method in ENMeval^25^ with an aggregation factor of 5. We selected the best model for each species based on sequentially considering the lowest measure of omission rate at the 10^th^ percentile (OR10) and the highest values of the Area Under the Curve (AUC) ^19,20,26^.

### Functional traits

Our target species represent a broad range of life-history strategies and climatic preferences, occurring in life zones ranging from tropical wet forests to subtropical and dry forests. Across the range of climatically suitable habitat for our study species, median precipitation in the driest month ranged from 1176 ±SD 271 mm to 2856 ±SD 849 mm (Table 1). We report data on functional traits expanded from previously published work ^27^. Specifically, we reported new data for wood density (g cm^-3^), leaf thickness (mm), and leaf specific area (SLA, m^2^ kg^-1^). Data collection for these traits followed the methods described in ^27^. Values for maximum height (m) were obtained from the species descriptions in Little and Wadsworth ^15^. Data on seed dry mass was compiled from the Seed Information Database (https://ser-sid.org/) and ^28,29^.

## DATA RECORD

All raw reads from the SLAF-seq sequencing are available at the European Nucleotide Archive (ENA, https://www.ebi.ac.uk/ena/browser/home) under the project accession number PRJEB96042. Output from species distribution models and new trait data are deposited at (https://zenodo.org/uploads/17279448).

## TECHNICAL VALIDATION

### Quality control, filtering, and SNP calling

We used FastQC v0.12.0 ^30^ (https://www.bioinformatics.babraham.ac.uk/projects/fastqc/) to check the quality of the raw reads of each sample and then combined the quality reports with multiQC v1.22.2 ^31^. We removed additional adapter content with fastp v0.23.4 ^32^ in pair-end mode (PE parameter, forward adapter: AGATCGGAAGAGCACACGTCTGAACTCCAGTCAC, reverse adaptor: AGATCGGAAGAGCGTCGTGTAGGGAAAGAGTGT). We unified the length of the reads with Trimmomatic v0.40 ^33^. We first removed any additional adaptor sequence with the option ILLUMINACLIP with the TruSeq3-PE-2.fa adapter file, allowing 2 seed mismatches, a palindrome clip threshold of 30, and a simple clip threshold of 10. We then cut the reads to a unique length of 120 bp and filtered out reads shorter than 120 bp using the following options CROP:130, HEADCROP:10, and MINLEN:120. For each species independently, we then used Stacks v2.66 ^34^ to align the reads to each other and identify SNPs using the ‘*de novo*’ pipeline (i.e., without reference genome) with default parameters. We optimized the number of mismatches between stacks by running the pipeline with -M parameter values ranging from 1 to 5. We chose M = 5 (i.e., maximum ∼1 SNP every 24 bp) as the most appropriate parameter as it led to the highest number of high-quality SNPs. In the ‘*de novo*’ pipeline, we kept only loci present in at least 80% of individuals in each population, using the parameter --min-samples-per-pop at 0.8.

In the next step, for each species, we independently performed SNP-level and sample-level filtering across all individuals. We first retained SNPs with <0.8 missing data and then removed samples with > 0.5 missing data. Finally, we used the ‘*rCNV*’ R package v1.3.9 ^35^) with default parameters to identify and filter out pseudo-SNPs stemming from putative paralogous and copy number variant (CNV) regions (Supplementary Figure 1). When ignored, population structure can affect the apparent excess of heterozygotes, a metric used to identify pseudo-SNPs. To control for this effect, we used the *F*_*IS*_ parameter in the *allele*.*info* function in the ‘*rCNV*’ R package ^35^.

### Population genetics statistics

We computed population genetics summary statistics using the Stacks v2.66^34^ program ‘Populations’. In order to evaluate the potential effect of linkage disequilibrium, we calculated the summary statistics with two datasets, one with all SNPs (using the **-**-vcf parameter in *‘Populations’*) and one with only one randomly selected SNP per SLAF-tag (using the **-**-write-random-snp parameter in *‘Populations’*). Due to the high correlation between statistics computed from either dataset (Pearson’s correlation coefficient *r* = 0.995 and *r* = 0.997 for *F*_*IS*_ and *H*_*o*_, respectively, Supplementary Figure 2), we performed all downstream analyses using the full SNP datasets. We computed summary statistics at both the ‘island-level’ (i.e., pooling all samples per island, Supplementary Table 2) and ‘population-level’ for Puerto Rican samples (Table 2).

**Table 2.** Summary statistics based on Puerto Rico samples. The number of SNPs obtained after filtering, inbreeding coefficient (*F*_*IS*_), observed heterozygosity (*H*_*o*_), and all sites Nei’s nucleotide diversity (*π*) relative to the total number of sequenced sites, i.e. including monomorphic sites. Values presented are means (± standard deviation) across populations (sample sites) in Puerto Rico only. See Supplementary Table 2 for island-level values.

| Species | Number of SNPs | Population (samples) | $H_o$ | $H_e$ | $F_{IS}$ | $\pi$ |
| --- | --- | --- | --- | --- | --- | --- |
| *Amyris elemifera* | 302,719 | 3 (10) | 0.089 ± 0.014 | 0.073 ± 0.005 | 0.006 ± 0.006 | 0.0018 |
| *Bursera simaruba* | 273,840 | 10 (50) | 0.106 ± 0.011 | 0.104 ± 0.009 | 0.024 ± 0.007 | 0.0043 |
| *Casearia arborea* | 166,697 | 5 (22) | 0.146 ± 0.011 | 0.132 ± 0.026 | 0.020 ± 0.009 | 0.0030 |
| *Casearia guianensis* | 187,411 | 5 (16) | 0.141 ± 0.002 | 0.124 ± 0.012 | 0.013 ± 0.006 | 0.0028 |
| *Casearia sylvestris* | 189,775 | 9 (42) | 0.083 ± 0.003 | 0.078 ± 0.004 | 0.014 ± 0.004 | 0.0024 |
| *Cecropia schreberiana* | 411,061 | 10 (50) | 0.077 ± 0.017 | 0.08 ± 0.021 | 0.027 ± 0.020 | 0.0068 |
| *Dacryodes excelsa* | 58,330 | 4 (16) | 0.126 ± 0.008 | 0.110 ± 0.006 | 0.014 ± 0.017 | 0.0015 |
| *Faramea occidentalis* | 80,061 | 1 (4) | 0.139 | 0.129 | 0.019 | 0.0032 |
| *Hirtella rugosa* | 158,984 | 5 (24) | 0.171 ± 0.018 | 0.160 ± 0.007 | 0.028 ± 0.034 | 0.0019 |
| *Gymnanthes lucida* | 49,606 | 3 (15) | 0.19 ± 0.031 | 0.192 ± 0.037 | 0.056 ± 0.030 | 0.0018 |
| *Krugiodendron ferreum* | 124,491 | 3 (15) | 0.152 ± 0.013 | 0.136 ± 0.004 | 0.001 ± 0.019 | 0.0020 |
| *Manilkara bidentata* | 189,702 | 2 (9) | 0.19 ± 0.002 | 0.179 ± 0.001 | 0.035 ± 0.001 | 0.0032 |
| *Micropholis guyanensis* | 118,159 | 6 (25) | 0.139 ± 0.014 | 0.122 ± 0.027 | 0.017 ± 0.012 | 0.0024 |
| *Myrcia deflexa* | 120,169 | 5 (19) | 0.105 ± 0.008 | 0.087 ± 0.015 | 0.003 ± 0.008 | 0.0017 |
| *Poitea florida* | 156,227 | 3 (12) | 0.134 ± 0.003 | 0.124 ± 0.017 | 0.031 ± 0.011 | 0.0046 |
| *Prestoea acuminata* | 52,588 | 5 (23) | 0.066 ± 0.002 | 0.060 ± 0.002 | 0.010 ± 0.002 | 0.0010 |
| *Didymopanax morototoni* | 433,637 | 7 (34) | 0.06 ± 0.004 | 0.058 ± 0.002 | 0.015 ± 0.005 | 0.0030 |
| *Sloanea berteriana* | 92,273 | 3 (11) | 0.176 ± 0.008 | 0.148 ± 0.014 | 0.013 ± 0.021 | 0.0009 |
| *Tabebuia heterophylla* | 24,413 | 11 (47) | 0.045 ± 0.003 | 0.042 ± 0.005 | 0.010 ± 0.004 | 0.0032 |

### SDM evaluation statistics

Across the selected SDMs, the values of omission rate at the 10^th^ percentile ranged from 0.07 to 0.18 (expected value=0.10; median 0.10 ± SD 0.03). Values for the Area Under the Curve (AUC) ranged from 0.60 to 0.85 (median 0.74 ± SD 0.07). The SDM performance metrics for all species are reported in Supplementary Table 3.

## DATA OVERVIEW

We computed several commonly-used population genetics statistics as a validation of genotyping quality. After filtering, the total number of samples across all species retained was 661, including 450 samples from Puerto Rico, 101 from Hispañola (Dominican Republic), and 110 from the US Virgin Islands. Across species, the number of species-specific single-nucleotide polymorphisms (SNPs) ranged from 24,413 to 433,637 SNPs. Island-wide summary statistics showed consistent patterns in that differences were more pronounced between species than among islands within species (Supplementary Table 2). Hereafter, we report population-level summary statistics focusing on the populations in Puerto Rico (Table 2, Figure 2). Overall, our estimates of genetic diversity are consistent with expectations for obligate outcrossing plant species. Across species, mean values of observed heterozygosity (*H*_*o*_) were rather low (ranging from 0.040 ±SD 0.003 to 0.190 ±SD 0.031) and were comparable to the levels of expected heterozygosity (*H*_*e*_; 0.040 ±SD 0.005 to 0.190 ±SD 0.037), indicating no notable patterns of outbreeding or inbreeding. All values of the inbreeding coefficient (*F*_*is*_) were close to 0 (ranging from 0.001 ±SD 0.019 to 0.056 ±SD 0.03). Species-level Nei’s nucleotide diversity computed across all SNPs and corrected for non-variant sites (*π*) ranged from 0.0009 to 0.0068 (Figure 2), in line with values reported in a previous study on lowland tropical trees ^11^. All species except one (*Cecropia schreberiana*) showed low variation across populations, which is also consistent with observations of temperate tree species ^36^.

**Figure 2.**
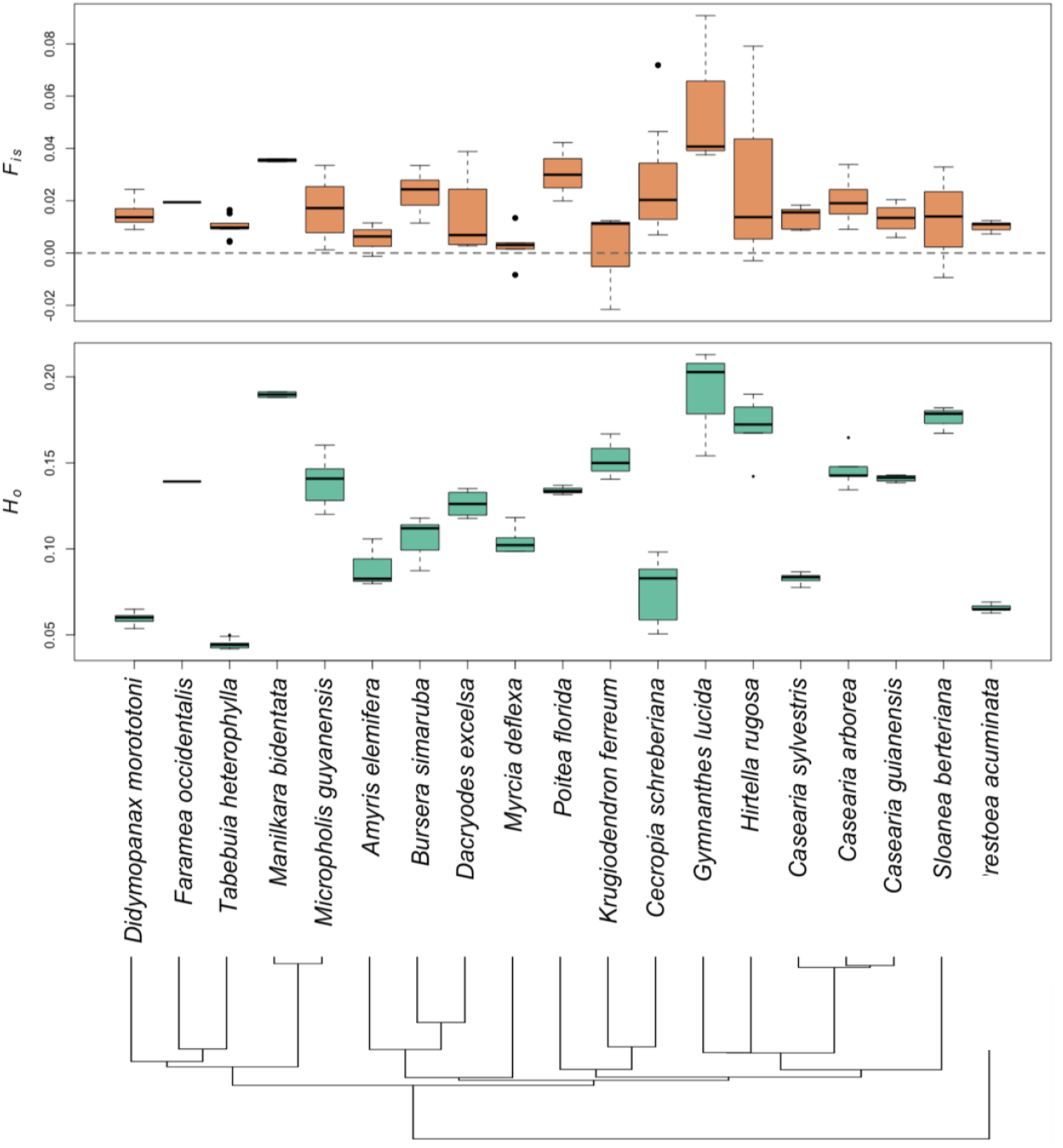
Box plots showing the values of the inbreeding coefficient (*F*_*IS*_) observed and heterozygosity (*H*_*o*_) computed for each population of each species in Puerto Rico. Species are arranged by phylogenetic relationships based on the island-wide phylogeny of ^37^. Island-level values for various statistics are provided in Supplementary Table 2.

## DATA AVAILABILITY

All raw reads from the SLAF-seq sequencing are available at the European Nucleotide Archive (ENA, https://www.ebi.ac.uk/ena/browser/home) under the project accession number PRJEB96042. Output from species distribution models and new trait data are deposited in Zenodo (https://zenodo.org/uploads/17279448)

## CODE AVAILABILITY

The costume scripts used to build the species distribution models is available in Zenodo (https://zenodo.org/uploads/17279448)

## ACKNOWLEDGEMENTS

We thank Brandon Ruiz for assistance with sample collection. Research was funded by grants from the Swedish Research Council, Formas (2020-00921 to RM and PM), the Swedish Research Council, Vetenskapsrådet (2019-03758 to RM), the Swedish Phytogeographical Society (to LM and RM), the Birgitta Sintring Foundation (to LM), and the US National Science Foundation (NSF DEB-1753810 to MU and RM). Additional support was provided by the Luquillo LTER (US NSF grant DEB-1831952), the University of Puerto Rico-Río Piedras, and the USDA Forest Service International Institute of Tropical Forestry.

## AUTHOR CONTRIBUTIONS

LM, PM, and RM designed the study. LM, BCG, TC, FBS, EG, JP and RM performed fieldwork. LM performed DNA extractions with support from FBS and PM. LM, MU and RM collected data on species distributions and functional traits. LM, PM, MU, and RM acquired funding. LM analyzed data with support from PM and RM. LM, PM, and RM wrote the initial draft, and all authors gave input towards the final version.

## FUNDING

Research was funded by grants from the Swedish Research Council, Formas (2020-00921 to RM and PM), the Swedish Research Council, Vetenskapsrådet (2019-03758 to RM), the Swedish Phytogeographical Society (to LM and RM), the Birgitta Sintring Foundation (to LM), and the US National Science Foundation (NSF DEB-1753810 to MU and RM)

## COMPETING INTERESTS

The authors have no competing interests to declare.

## Notes

https://www.ebi.ac.uk/ena/browser/home

https://zenodo.org/uploads/17279448

### Competing Interest Statement

The authors have declared no competing interest.

