## Supplementary material for "A comparative dataset on population genetics, traits, and distributions for nineteen Caribbean tree species": Suplementary information

### Table of contents

1. **Supplementary Figure 1:** Detection of pseudo-SNPs
2. **Supplementary Figure 2:** Correlation between population genetic estimates when computed from all SNPs or one SNP per tag
3. **Supplementary Table 1:** Species-level extractions quality control
4. **Supplementary Table 2:** Island-level population genetics summary statistics
5. **Supplementary Table 3:** SDM performance statistics

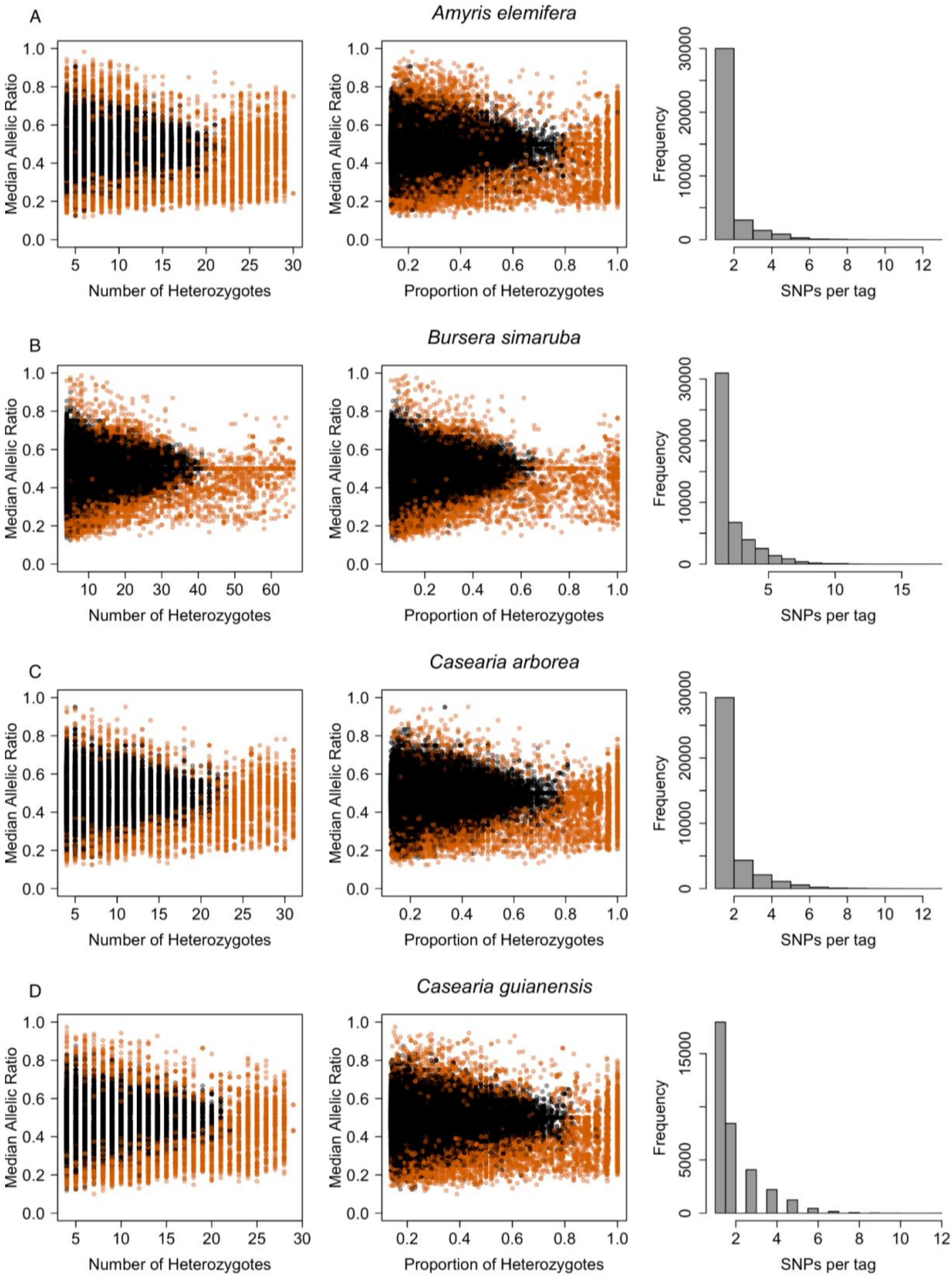
A
Amyris elemifera
Median Allelic Ratio
Number of Heterozygotes
Proportion of Heterozygotes
Frequency
SNPs per tag
B
Bursera simaruba
Median Allelic Ratio
Number of Heterozygotes
Proportion of Heterozygotes
Frequency
SNPs per tag
C
Casearia arborea
Median Allelic Ratio
Number of Heterozygotes
Proportion of Heterozygotes
Frequency
SNPs per tag
D
Casearia guianensis
Median Allelic Ratio
Number of Heterozygotes
Proportion of Heterozygotes
Frequency
SNPs per tag

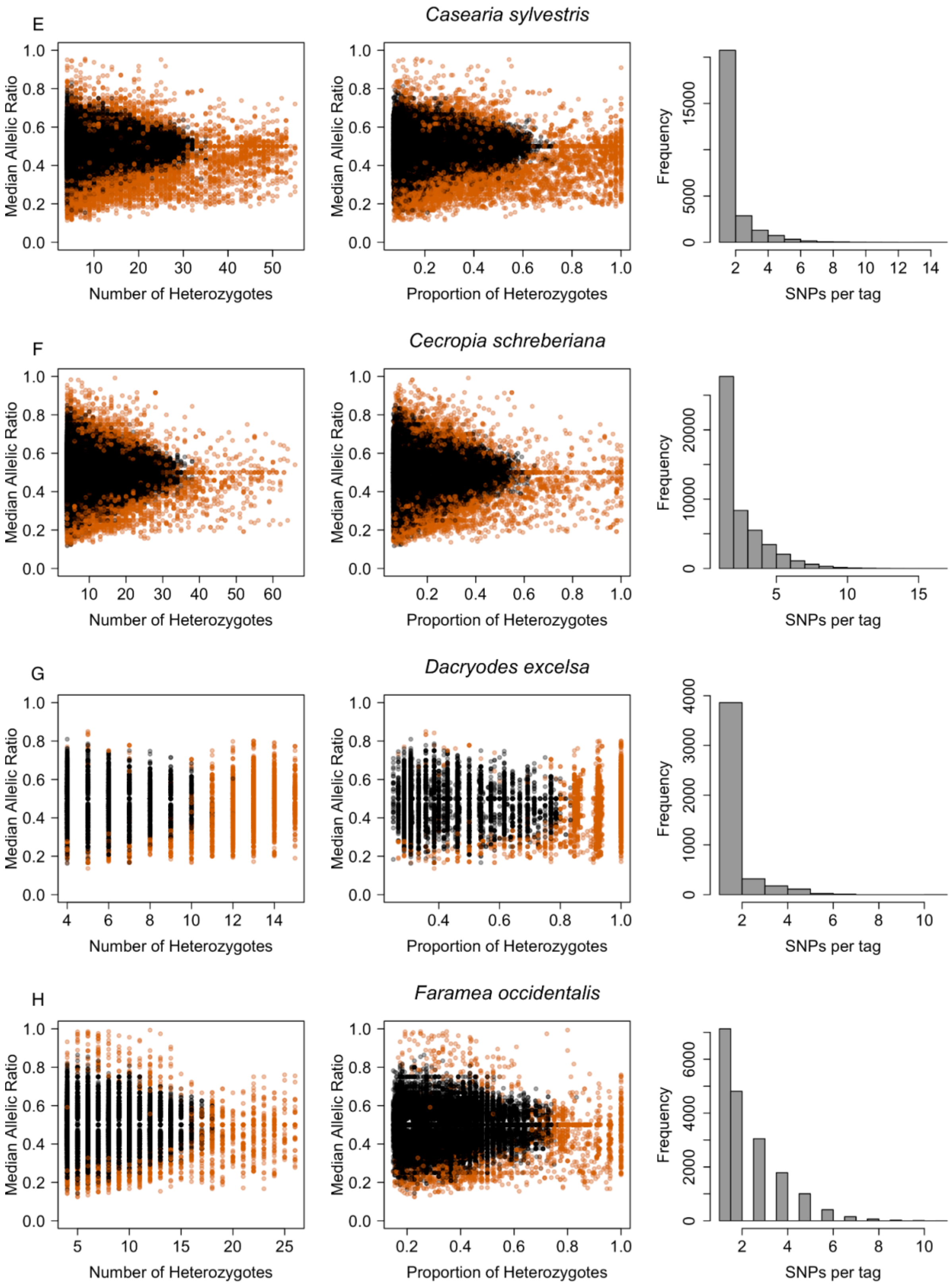

E
Casearia sylvestris
F
Cecropia schreberiana
G
Dacryodes excelsa
H
Faramea occidentalis
Median Allelic Ratio
Number of Heterozygotes
Proportion of Heterozygotes
Frequency
SNPs per tag

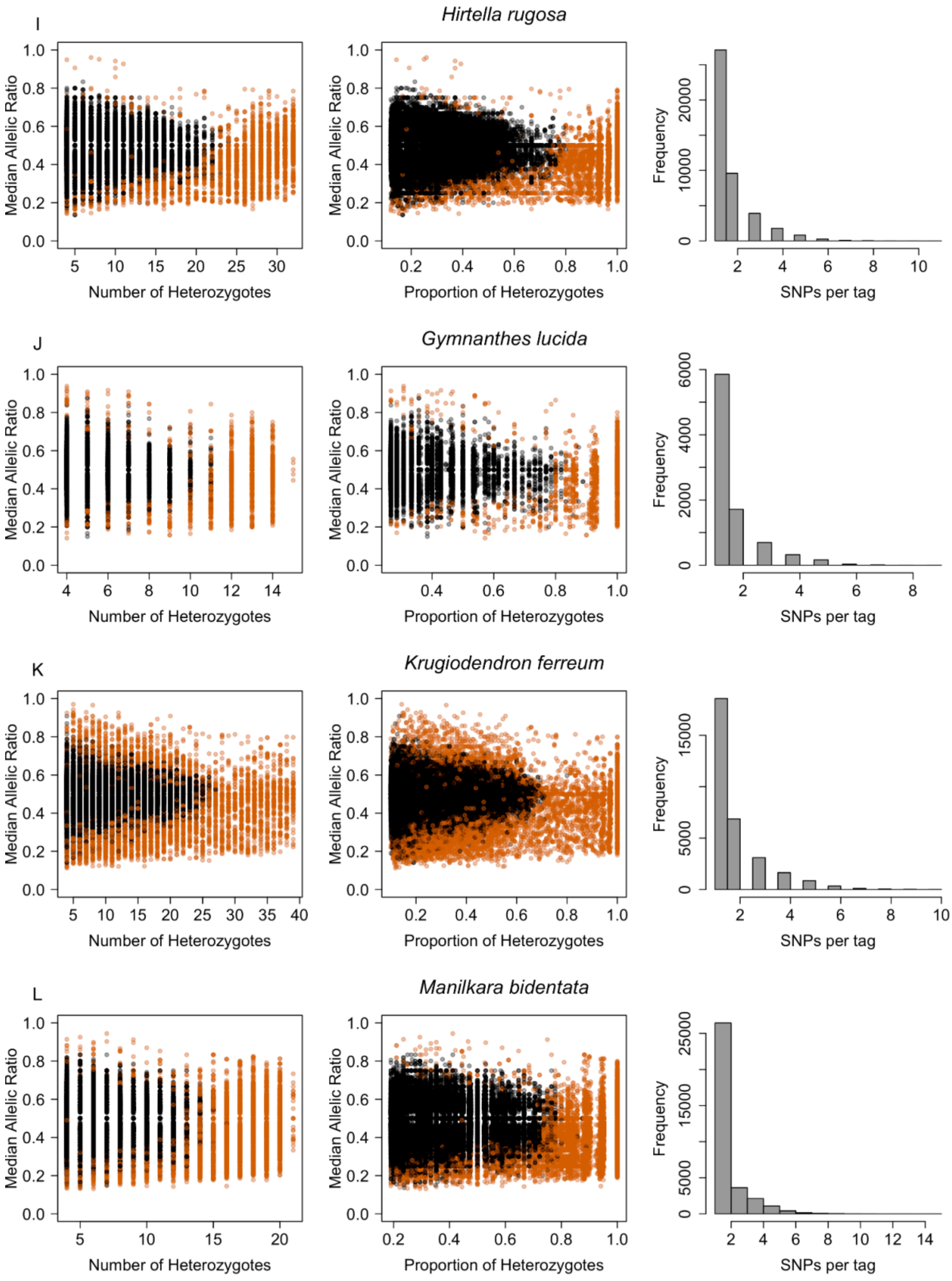
I
Hirtella rugosa
Median Allelic Ratio
Number of Heterozygotes
Median Allelic Ratio
Proportion of Heterozygotes
Frequency
SNPs per tag
J
Gymnanthes lucida
Median Allelic Ratio
Number of Heterozygotes
Median Allelic Ratio
Proportion of Heterozygotes
Frequency
SNPs per tag
K
Krugiodendron ferreum
Median Allelic Ratio
Number of Heterozygotes
Median Allelic Ratio
Proportion of Heterozygotes
Frequency
SNPs per tag
L
Manilkara bidentata
Median Allelic Ratio
Number of Heterozygotes
Median Allelic Ratio
Proportion of Heterozygotes
Frequency
SNPs per tag

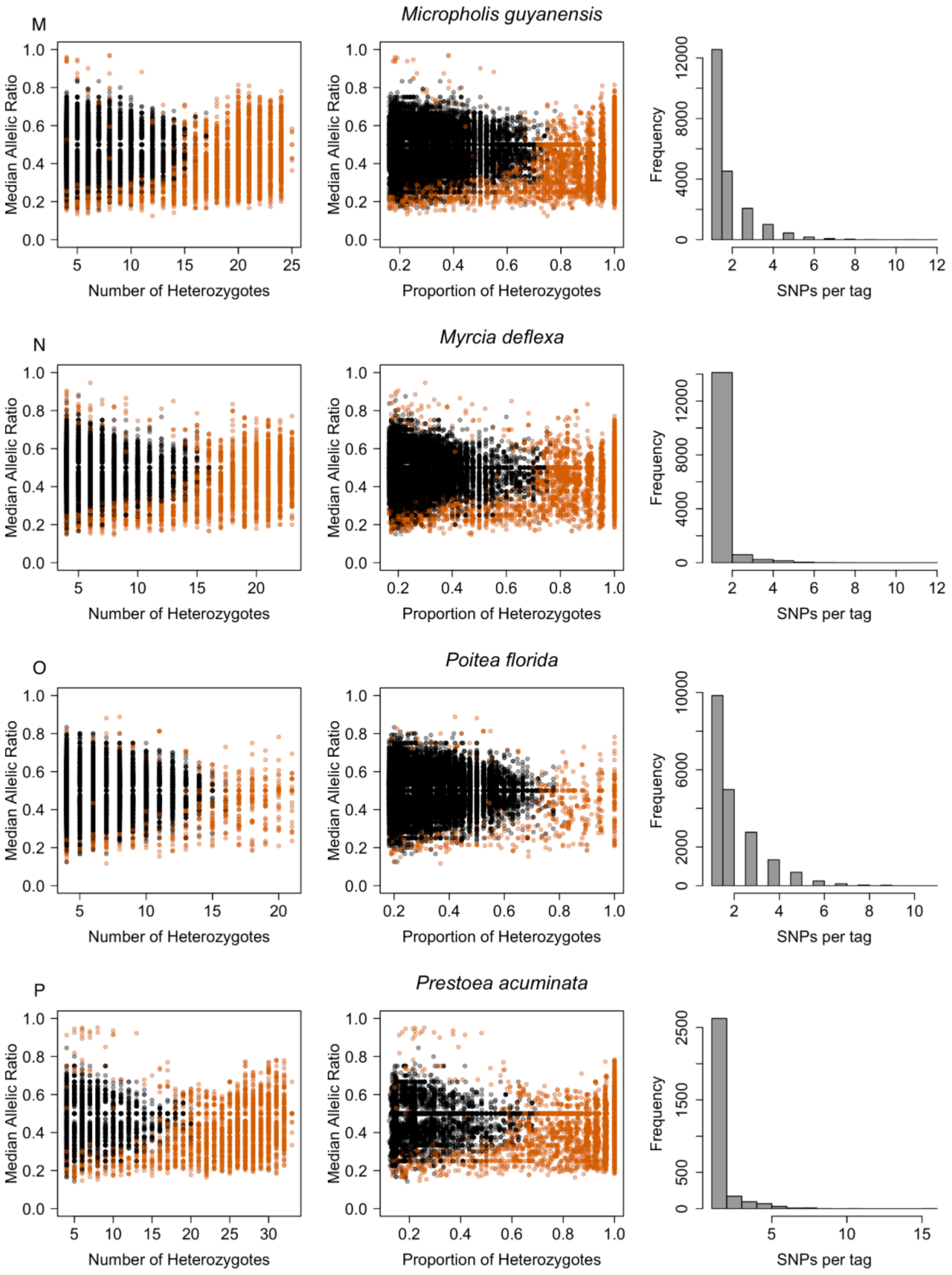
M
Micropholis guyanensis
Median Allelic Ratio
Number of Heterozygotes
Proportion of Heterozygotes
Frequency
SNPs per tag
N
Myrcia deflexa
Median Allelic Ratio
Number of Heterozygotes
Proportion of Heterozygotes
Frequency
SNPs per tag
O
Poitea florida
Median Allelic Ratio
Number of Heterozygotes
Proportion of Heterozygotes
Frequency
SNPs per tag
P
Prestoea acuminata
Median Allelic Ratio
Number of Heterozygotes
Proportion of Heterozygotes
Frequency
SNPs per tag

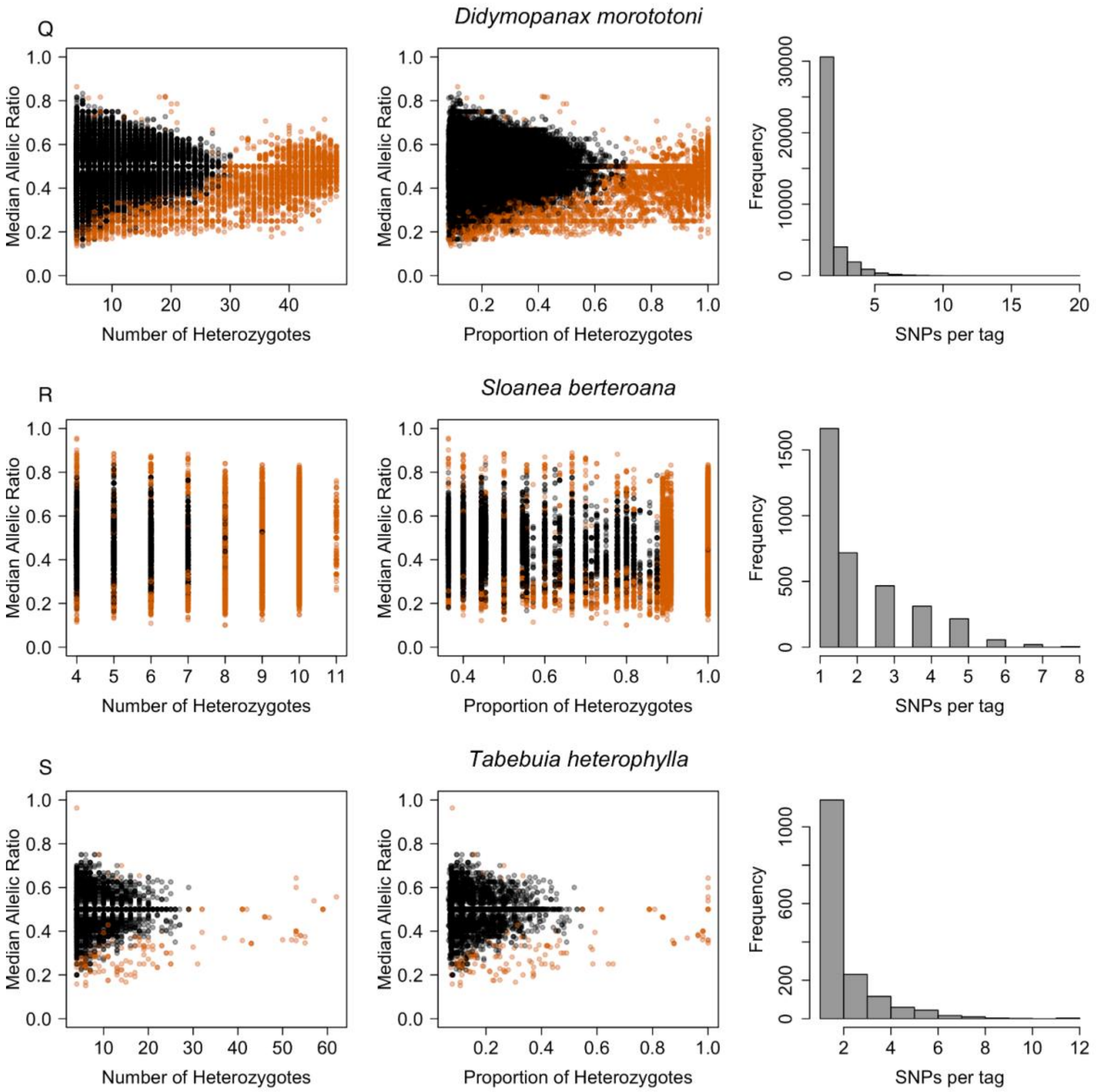


**Supplementary Figure 1**: **Detection of pseudo-SNPs.** For each species (rows), the left and middle panels display SNP-based median allelic depth of coverage ratios as function of the number of heterozygotes and their proportion, respectively. Pseudo-SNPs (orange) were identified and excluded from the dataset using the *rCNV* R package (v1.3.9 [36]). The right panel is the distribution of the number of SNPs per tag after pruning for pseudo-SNPs.

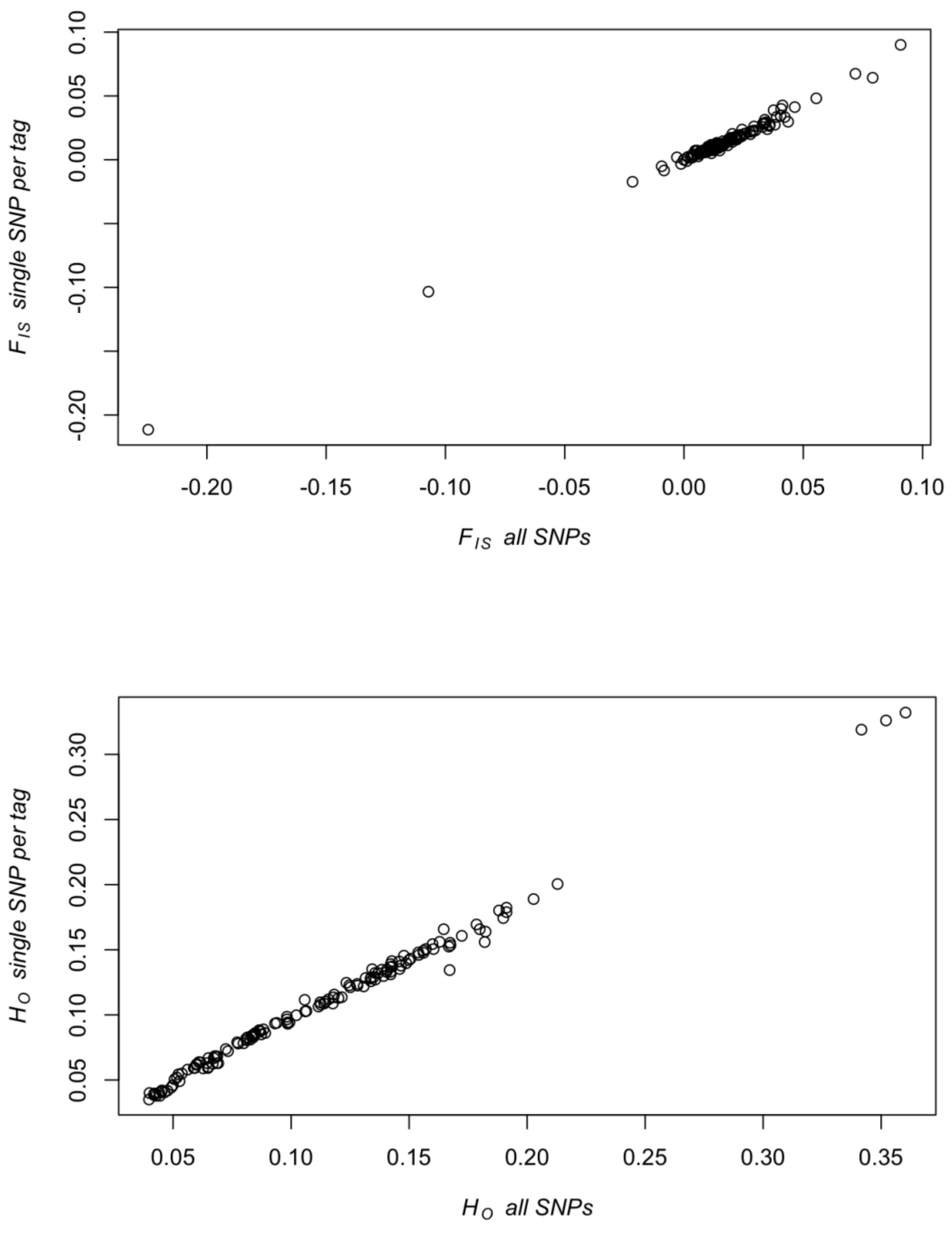


**Supplementary Figure 2:** Correlation between population genetic estimates when computed from all SNPs or one SNP per tag. The top panel shows the correlation between $F_{IS}$ calculated across all SNPs and $F_{IS}$ calculated using one SNP per tag. The bottom panel shows the correlation between $H_0$ calculated across all SNPs and $H_0$ calculated using one SNP per tag.

**Supplementary Table 1.** Species-level quality and purity control mean values (Nanodrop) and concentration mean values (Qubit) for DNA extractions.

| Species | Concentration ug/ml (100 ul) | 260/280 | 230/260 |
|---|---|---|---|
| *Amyris elemifera* | 42.08 ± 29.09 | 1.75 ± 0.08 | 1.27 ± 0.32 |
| *Bursera simaruba* | 47.4 ± 17.57 | 1.79 ± 0.17 | 1.15 ± 0.34 |
| *Casearia arborea* | 34.13 ± 17.88 | 1.75 ± 0.1 | 1.23 ± 0.36 |
| *Casearia guianensis* | 37.01 ± 14.08 | 1.72 ± 0.11 | 1.17 ± 0.33 |
| *Casearia sylvestris* | 36.64 ± 13.64 | 1.6 ± 0.13 | 4.06 ± 16.15 |
| *Cecropia schreberiana* | 44.32 ± 16.63 | 1.76 ± 0.05 | 1.3 ± 0.27 |
| *Dacryodes excelsa* | 28.35 ± 11.13 | 1.56 ± 0.13 | 1.32 ± 0.26 |
| *Didymopanax morototoni* | 38.03 ± 21.91 | 1.69 ± 0.17 | 0.98 ± 0.34 |
| *Faramea occidentalis* | 30.74 ± 19.56 | 1.97 ± 2.2 | 1.43 ± 0.39 |
| *Gymnanthes lucida* | 29.47 ± 13.8 | 1.96 ± 0.09 | 1.32 ± 0.24 |
| *Hirtella rugosa* | 48.2 ± 9.46 | 2.85 ± 7.04 | 1.15 ± 0.36 |
| *Krugiodendron ferreum* | 34.6 ± 11.32 | 1.7 ± 0.15 | 1.17 ± 0.41 |
| *Manilkara bidentata* | 45.07 ± 21.58 | 1.75 ± 0.16 | 1.17 ± 0.23 |
| *Micropholis guyanensis* | 43.2 ± 13.38 | 1.72 ± 0.12 | 1.09 ± 0.3 |
| *Myrcia deflexa* | 45.23 ± 15.37 | 1.74 ± 0.07 | 1.43 ± 0.4 |
| *Poitea florida* | 34.8 ± 16.6 | 1.61 ± 0.09 | 0.94 ± 0.21 |
| *Prestoea acuminata* | 109.59 ± 45.39 | 1.64 ± 0.13 | 1.28 ± 0.3 |
| *Sloanea berteroana* | 44.96 ± 30.16 | 1.67 ± 0.1 | 0.74 ± 0.13 |
| *Tabebuia heterophylla* | 4.77 ± 3.2 | 1.83 ± 0.08 | 1.72 ± 0.36 |

**Supplementary Table 2.** Island-level population genetics summary statistics. For each species and each island the table gives the observed heterozygosity ($H_o$), all sites Nei's nucleotide diversity (pi), and the inbreeding coefficient ($F_{IS}$) either computed across all of the samples in each island ($H_o$ all, $F_{IS}$ all, Pi all), or per sampling location and we report the mean value with associated standard error.

| species | Island | $H_O$ | $H_O$ mean (± sd) | $F_{IS}$ | $F_{IS}$ mean (± sd) | $\pi$ | mean (± sd) |
|---|---|---|---|---|---|---|---|
| *Amyris elemifera* | PR | 0.0857 | 0.0894 ± 0.0082 | 0.0319 | 0.0055 ± 0.0037 | 0.0018 | 0.0017 ± 1e-04 |
| | DR | 0.0983 | - | 0.0173 | - | 0.0019 | - |
| | VI | 0.0892 | 0.0904 ± 0.0035 | 0.0213 | 0.0124 ± 0.0018 | 0.0017 | 0.0017 ± 1e-04 |
| *Bursera simaruba* | PR | 0.1063 | 0.1074 ± 0.0033 | 0.1024 | 0.0214 ± 0.0029 | 0.0043 | 0.0038 ± 1e-04 |
| | DR | 0.1209 | 0.1219 ± 0.0061 | 0.0599 | 0.0346 ± 0.0059 | 0.0048 | 0.0046 ± 1e-04 |
| | VI | 0.0657 | 0.0658 ± 0.0015 | 0.0213 | 0.0116 ± 5e-04 | 0.0024 | 0.0023 ± 1e-04 |
| *Casearia arborea* | PR | 0.1481 | 0.1463 ± 0.0051 | 0.0726 | 0.0202 ± 0.0042 | 0.003 | 0.0027 ± 1e-04 |
| | DR | 0.1493 | 0.1492 ± 0.0137 | 0.1177 | 0.0383 ± 0.0171 | 0.0034 | 0.0029 ± 4e-04 |
| *Casearia guianensis* | PR | 0.1406 | 0.1396 ± 0.0016 | 0.0343 | 0.0106 ± 0.0035 | 0.0028 | 0.0027 ± 1e-04 |
| | DR | 0.1585 | 0.1585 ± 0.0014 | 0.0405 | 0.0218 ± 0.0015 | 0.0032 | 0.0031 ± 0 |
| | VI | 0.1234 | - | 0.0157 | - | 0.0024 | - |
| *Casearia sylvestris* | PR | 0.0828 | 0.083 ± 9e-04 | 0.0538 | 0.0141 ± 0.0014 | 0.0024 | 0.0023 ± 0 |
| | DR | 0.0732 | - | 0.0164 | - | 0.002 | - |
| | VI | 0.0687 | - | 0.0075 | - | 0.0019 | - |
| *Cecropia schreberia* | PR | 0.0766 | 0.077 ± 0.0055 | 0.1168 | 0.0267 ± 0.0064 | 0.0068 | 0.0051 ± 4e-04 |
| | DR | 0.0731 | 0.0716 ± 0.0123 | 0.0344 | 0.0148 ± 0.0042 | 0.005 | 0.0045 ± 8e-04 |
| | VI | 0.0682 | | 0.0167 | - | 0.0043 | 0.0043 ± |
| *Dacryodes excelsa* | PR | 0.1263 | 0.1262 ± 0.004 | 0.056 | 0.0138 ± 0.0085 | 0.0015 | 0.0015 ± 1e-04 |
| *Faramea occidentalis* | PR | 0.2128 | 0.2462 ± 0.0566 | 0.0938 | -0.0338 ± 0.0434 | 0.0055 | 0.0049 ± 0.001 |
| | DR | 0.3403 | 0.3361 ± 0.0051 | -<br>0.2212 | -0.1057 ± 0.062 | 0.0047 | 0.0059 ± 6e-04 |
| | VI | 0.1322 | 0.1319 ± 0.004 | 0.0353 | 0.0219 ± 6e-04 | 0.0031 | 0.0031 ± 1e-04 |

| | | | | | | | |
|---|---|---|---|---|---|---|---|
| ***Gymnathes lucida*** | PR | 0.1683 | 0.1708 ± 0.0082 | 0.0624 | 0.0278 ± 0.0151 | 0.0019 | 0.0018 ± 0 |
| | VI | 0.1742 | 0.1737 ± 0.0062 | 0.0475 | 0.0136 ± 0.0084 | 0.0019 | 0.0018 ± 1e-04 |
| ***Hirtella rugosa*** | PR | 0.1893 | 0.1899 ± 0.0181 | 0.2563 | 0.0564 ± 0.0172 | 0.0018 | 0.0013 ± 1e-04 |
| ***Krugiodendron ferreum*** | PR | 0.1521 | 0.1524 ± 0.0077 | 0.0484 | 7e-04 ± 0.0111 | 0.002 | 0.0018 ± 0 |
| | DR | 0.1511 | 0.1513 ± 0.0025 | 0.0452 | 0.023 ± 0.0127 | 0.002 | 0.0019 ± 1e-04 |
| | VI | 0.1524 | 0.1526 ± 0.0034 | 0.0276 | 0.0118 ± 0.0043 | 0.0019 | 0.0018 ± 0 |
| ***Manilkara bidentata*** | PR | 0.1892 | 0.1897 ± 0.0016 | 0.0595 | 0.0355 ± 5e-04 | 0.0032 | 0.0031 ± 0 |
| | DR | 0.1506 | - | 0.0381 | - | 0.0025 | - |
| | VI | 0.1913 | - | 0.0211 | - | 0.0031 | - |
| ***Micropholis guyanensis*** | PR | 0.1434 | 0.1395 ± 0.0059 | 0.0961 | 0.017 ± 0.0048 | 0.0024 | 0.0021 ± 1e-04 |
| ***Myrcia deflexa*** | PR | 0.1046 | 0.1048 ± 0.0037 | 0.0597 | 0.0027 ± 0.0035 | 0.0017 | 0.0015 ± 1e-04 |
| | DR | 0.0685 | - | 0.0413 | - | 0.0012 | - |
| ***Poitea florida*** | PR | 0.1338 | 0.134 ± 0.0016 | 0.0875 | 0.0307 ± 0.0064 | 0.0046 | 0.0041 ± 1e-04 |
| | VI | 0.1249 | 0.1249 ± 2e-04 | 0.0431 | 0.0315 ± 0.0025 | 0.0039 | 0.0039 ± 0 |
| ***Prestoea acuminatata*** | PR | 0.0648 | 0.0657 ± 0.0011 | 0.0539 | 0.0102 ± 9e-04 | 0.001 | 0.0009 ± 0 |
| | DR | 0.0527 | - | 0.0149 | - | 0.0007 | - |
| ***Didymopanax morototoni*** | PR | 0.0597 | 0.0596 ± 0.0014 | 0.0381 | 0.0149 ± 0.002 | 0.003 | 0.0029 ± 0 |
| | DR | 0.0699 | 0.07 ± 0.0024 | 0.0197 | 0.0096 ± 0.0018 | 0.0033 | 0.0032 ± 1e-04 |
| | VI | 0.0522 | - | 0.0101 | - | 0.0024 | - |
| ***Sloanea berteroana*** | PR | 0.176 | 0.1759 ± 0.0045 | 0.0578 | 0.0125 ± 0.0122 | 0.0009 | 0.0009 ± 1e-04 |
| ***Tabebuia heterophylla*** | PR | 0.0444 | 0.0449 ± 8e-04 | 0.0738 | 0.0092 ± 0.0014 | 0.0032 | 0.0026 ± 1e-04 |
| | VI | 0.042 | 0.0423 ± 0.0018 | 0.0369 | 0.0093 ± 0.0019 | 0.0028 | 0.0025 ± 1e-04 |

**Supplementary Table 3:** SDM performance statistics.

| Species | fc | rm | auc.train | cbi.train | auc.diff ± sd | auc.val ± sd | cbi.val ± sd | or.10p ± sd | or.mtp ± sd | delta.AICc |
|---|---|---|---|---|---|---|---|---|---|---|
| *Amyris elemifera* | H | 2 | 0.79 | 0.89 | 0.04 ± 0.04 | 0.76 ± 0.04 | 0.32 ± 0.5 | 0.1 ± 0.09 | 0.04 ± 0.05 | 4.08 |
| *Bursera simaruba* | L | 2 | 0.74 | 0.93 | 0.03 ± 0.01 | 0.72 ± 0.01 | 0.6 ± 0.24 | 0.11 ± 0.07 | 0.03 ± 0.05 | 4.27 |
| *Casearia arborea* | LQH | 3 | 0.79 | 0.9 | 0.05 ± 0.03 | 0.77 ± 0.04 | 0.63 ± 0.11 | 0.1 ± 0.07 | 0.02 ± 0.02 | 0.96 |
| *Casearia guianensis* | LQ | 5 | 0.73 | 0.94 | 0.02 ± 0.01 | 0.72 ± 0.02 | 0.72 ± 0.12 | 0.09 ± 0.07 | 0.01 ± 0.02 | 9.42 |
| *Casearia sylvestris* | H | 3 | 0.69 | 0.95 | 0.03 ± 0.01 | 0.68 ± 0.03 | 0.64 ± 0.23 | 0.11 ± 0.03 | 0.01 ± 0.02 | 7.89 |
| *Cecropia schreberiana* | LQH | 4 | 0.75 | 0.97 | 0.03 ± 0.01 | 0.73 ± 0.03 | 0.85 ± 0.09 | 0.1 ± 0.05 | 0.01 ± 0.01 | 0 |
| *Dacryodes excelsa* | H | 2 | 0.85 | 0.9 | 0.07 ± 0.07 | 0.79 ± 0.05 | 0.51 ± 0.38 | 0.15 ± 0.13 | 0.13 ± 0.13 | 2.64 |
| *Faramea occidentalis* | LQH | 3 | 0.72 | 0.87 | 0.07 ± 0.06 | 0.65 ± 0.06 | 0.26 ± 0.21 | 0.16 ± 0.13 | 0.03 ± 0.05 | 0 |
| *Gymnanthes lucida* | LQ | 5 | 0.76 | 0.77 | 0.05 ± 0.03 | 0.75 ± 0.05 | 0.45 ± 0.28 | 0.09 ± 0.1 | 0.03 ± 0.05 | 12.28 |
| *Hirtella rugosa* | L | 4 | 0.78 | 0.9 | 0.11 ± 0.08 | 0.74 ± 0.12 | 0.59 ± 0.55 | 0.18 ± 0.27 | 0.04 ± 0.08 | 1.02 |
| *Krugiodendron ferreum* | L | 1 | 0.83 | 0.82 | 0.06 ± 0.06 | 0.78 ± 0.05 | 0.6 ± 0.12 | 0.19 ± 0.07 | 0.08 ± 0.06 | 0 |
| *Manilkara bidentata* | LQH | 2 | 0.82 | 0.88 | 0.08 ± 0.08 | 0.76 ± 0.08 | 0.52 ± 0.24 | 0.18 ± 0.18 | 0.07 ± 0.09 | 6.79 |
| *Micropholis guyanensis* | LQH | 4 | 0.87 | 0.84 | 0.05 ± 0.02 | 0.86 ± 0.04 | 0.8 ± 0.09 | 0.11 ± 0.11 | 0.04 ± 0.08 | 9.53 |
| *Myrcia deflexa* | L | 4 | 0.75 | 0.93 | 0.07 ± 0.05 | 0.73 ± 0.07 | 0.64 ± 0.16 | 0.12 ± 0.11 | 0.02 ± 0.03 | 21.74 |
| *Poitea florida* | H | 5 | 0.68 | 0.9 | 0.13 ± 0.12 | 0.6 ± 0.12 | 0.33 ± 0.6 | 0.13 ± 0.15 | 0.11 ± 0.16 | 6.23 |
| *Prestoea acuminata* | L | 5 | 0.84 | 0.83 | 0.07 ± 0.04 | 0.83 ± 0.06 | 0.7 ± 0.12 | 0.08 ± 0.1 | 0.05 ± 0.1 | 4.17 |
| *Didymopanax morototoni* | H | 5 | 0.78 | 0.9 | 0.04 ± 0.02 | 0.75 ± 0.02 | 0.6 ± 0.13 | 0.15 ± 0.06 | 0.04 ± 0.03 | 3.73 |
| *Sloanea berteroana* | H | 2 | 0.88 | 0.94 | 0.06 ± 0.05 | 0.85 ± 0.07 | 0.4 ± 0.43 | 0.08 ± 0.09 | 0.06 ± 0.08 | 0 |
| *Tabebuia heterophylla* | L | 2 | 0.62 | 0.91 | 0.04 ± 0.03 | 0.61 ± 0.04 | 0.59 ± 0.22 | 0.09 ± 0.05 | 0.01 ± 0.02 | 11.98 |
